# Structural and functional dissection of the interplay between lipid and Notch binding by human Notch ligands

**DOI:** 10.1101/101394

**Authors:** Richard J. Suckling, Boguslawa Korona, Patricia Whiteman, Chandramouli Chillakuri, Laurie Holt, Penny A. Handford, Susan M. Lea

## Abstract

Recent data have expanded our understanding of Notch signaling by identifying a C2 domain at the N-terminus of Notch ligands which has both lipid- and receptor-binding properties. We present novel structures of human ligands Jagged2 and DLL4 and human Notch-2, together with functional assays, which suggest that ligand-mediated coupling of membrane recognition and Notch binding is likely to be critical in establishing the optimal context for Notch signaling. Comparisons between the Jagged and Delta family show a huge diversity in the structures of the loops at the apex of the C2 domain implicated in membrane recognition and Jagged1 missense mutations which affect these loops and are associated with extrahepatic biliary atresia lead to a loss of membrane recognition, but do not alter Notch binding. Taken together, these data suggest that C2 domain binding to membranes is an important element in tuning ligand-dependent Notch signaling in different physiological contexts.

## Introduction

The Notch signaling pathway is conserved across all metazoan species, and plays key roles in many aspects of cell biology including cell-fate determination, stem cell maintenance, immune system activation and angiogenesis in humans (Bray, 2016; Guruharsha et al., 2012). Aberrant Notch signaling results in a number of inherited and acquired disorders, including various cancers, and it is therefore a key target for therapeutic intervention (Groth and Fortini, 2012; Hansson et al., 2004). Both the Notch receptors and the ligands are single-pass type I transmembrane proteins and direct protein-protein contact between adjacent cells initiates an intracellular signaling pathway. The Notch receptor exists as a heterodimeric transmembrane protein with the N-terminal extracellular domain consisting of up to 36 tandem epidermal growth factor-like (EGF) repeats. Binding of a Notch ligand to EGF11-12 of a Notch receptor results in a series of proteolytic cleavages, with the final intra-membrane cleavage by gamma secretase, causing the release of the intracellular domain of Notch (NICD) from the plasma membrane. Once released NICD translocates to the nucleus where it binds to a transcription factor of the CBF1, Suppressor of Hairless, Lag-1 (CSL) family in complex with the coactivator MAML. This complex then relieves repression and activates Notch target genes of the *Hes* and *Hey* repressor families (Nam et al., 2006). Whilst *Drosophila* have one Notch receptor, mammalian species have four (Notch1-4). Notch-ligand interactions can result in activation or inhibition of Notch signaling, depending on whether ligands are presented to Notch on neighbouring cells (*trans*), or on the same cell (*cis*) (Sprinzak et al., 2010).

There are four canonical cell surface mammalian Notch ligands, Jagged1, Jagged2, Delta-like1 (DLL1), and Delta-like 4 (DLL4), and one non-canonical ligand, DLL3, which is unable to activate Notch and is found predominantly in the Golgi apparatus (Geffers et al., 2007; Ladi et al., 2005; Serth et al., 2015). All of the Notch ligands have a modular extracellular architecture consisting of an N-terminal C2 domain (formerly known as the MNNL domain), a Delta/Serrate/Lag-2 (DSL) domain, and either 16 (Jagged1/Jagged2), 8 (DLL1/DLL4) or 7 (DLL3) EGF repeats. We have previously shown that the very N-terminal domain of human Jagged1 is a C2 domain, and lipid binding of this domain is required for optimal Notch activation (Chillakuri et al., 2013). This domain is conserved across the Notch ligands, however the loops within the putative lipid-binding site vary considerably between ligands, suggesting that the ligands have different lipid-binding specificity. Recently a co-crystal structure of a DLL4 variant N-EGF1 in complex with Notch1 EGF11-13 (Luca et al., 2015) has shown that Notch ligands interact via a platform located on one side of the N-terminal C2 domain away from the lipid binding region (site 1) and via their DSL domain (site 2), with Notch receptor domains EGF11 (site 2) and 12 (site 1) in an antiparallel fashion. This confirmed previous data showing that residues in the Jagged1 DSL domain, and in Notch1 EGF12 are critical for receptor-ligand interactions (Cordle et al., 2008; Whiteman et al., 2013). O-glycosylation of Notch plays an important role in regulating Notch signaling, with O-fucosylation on Thr-466 in EGF12 of Notch1 enhancing ligand binding (Stahl et al., 2008; Yao et al., 2011). In addition we have also shown that Fringe-catalysed addition of GlcNAc to the O-fucose at Thr-466 in EGF12 increases binding to ligands (Taylor et al., 2014). The DLL4-Notch1 complex structure shows that the O-fucose modification directly contributes to the binding interface, in addition to specific amino acid contacts (Luca et al., 2015).

Here we further highlight the variability of the C2 domain putative lipid-binding site in Notch ligands, by solving the crystal structures of N-terminal fragments of both human DLL4 and Jagged2. These new structures, together with a structure for Notch2 EGF11-13, have allowed a detailed comparison of both ligand/receptor and ligand/lipid-binding interactions. We further demonstrate *in vitro* that Notch receptor binding to ligand enhances interactions with lipids, suggesting that a ternary complex between Notch, ligand and lipid fine tunes generation of the Notch signal at the cell surface. A subset of *Jagged1* mutations which are associated with extrahepatic biliary atresia and affect the loops at the apex of the C2 domain reduce both Notch activation and lipid binding indicating the importance of membrane binding for tuning the Notch signal in specific physiological contexts.

## Results

### Structures of the N-terminal domains of human Notch ligands highlight conformational flexibility between EGF2-3 which may facilitate formation of an extended Notch/ligand binding interface

We expressed and solved the structures of various N-terminal fragments of both human DLL4 and Jagged2 (Table 1). These fragments include the N-terminal C2 lipid binding domain, the receptor binding DSL domain, and two (‘N-EGF2’) or three (‘N-EGF3’) adjacent EGF domains. Here we present the first structures of Jagged2 (N-EGF2 and N-EGF3) which has only 58% sequence identity with human Jagged1 (N-EGF3), together with the longest known structure of a DLL4 ligand (human DLL4 N-EGF3) (Figure 1A). This allows, for the first time, comparative structural analyses of all the canonical ligands. Superposition of the different Jagged2 structures from our study, with all the various Notch ligand structures (Chillakuri et al., 2013; Kershaw et al., 2015) across the DSL domain, shows that within this region the ligands form a near-linear domain organization (Figure 1B). However the angle between adjacent domains can vary subtly, resulting in the ligand structures appearing to fall into two groups, one group including Jagged1, DLL1 and some structures of Jagged2, and the other including DLL4, and some structures of Jagged2. The observation that Jagged2 is split across the two groups, suggests that there is some flexibility between adjacent domains of the Notch ligands, and that all may be able to adopt these different conformations. It is therefore likely that the conformation seen is determined by crystal packing. For example the angle between EGF2 and EGF3 in our hDLL4 structure, is likely due to interactions with a neighbouring molecule in the crystal lattice. The only ligand-Notch complex seen to date shows the ligand to be in the more bent conformation, we propose that this is due to the use of a short Notch EGF11-13 construct. We have previously shown that modelling of Notch EGF4-13 in complex with ligand, based on the published structure of the receptor/ligand complex, leads to a steric clash (Weisshuhn et al., 2016). However, if the ligand remodels into the straighter form, shown in our new structures, this allows good packing interactions between Notch EGF4-13 and ligand, extending the binding interface along the longitudinal axis, and suggests that remodelling is a prerequisite for the ligand to form optimal contacts with Notch (Figure 1C).

**Table 1.** Data Collection and refinement statistics

|  | Human Jagged2 |  |  | DLL4 N-EGF3 | Notch2 EGF11-13 |
| --- | --- | --- | --- | --- | --- |
|  | Jagged2 N-EGF2 | Jagged2 N-EGF2 | Apo-Jagged2 N-EGF3 |  |  |
| **Data Collection** |  |  |  |  |  |
| Beamline | Diamond I02 | Diamond I04-1 | Diamond I04-1 | Diamond I04-1 | Diamond I04 |
| Space group | P2<sub>1</sub>2<sub>1</sub>2<sub>1</sub> | P1 | P2<sub>1</sub>2<sub>1</sub>2<sub>1</sub> | C2 | P2<sub>1</sub>2<sub>1</sub>2<sub>1</sub> |
| Wavelength (Å) | 1.3 | 0.93 | 0.92 | 0.92 | 0.98 |
| **Cell dimensions (Å)** |  |  |  |  |  |
| a, b, c (Å) | 48.3, 83.9, 99.4 | 62.6, 92.9, 97.0 | 46.8, 77.2, 96.4 | 127.9, 49.8, 70.0 | 20.2, 49.8, 125.5 |
| α, β, γ (°) | 90.0, 90.0, 90.0 | 71.7, 83.2, 82.7 | 90.0, 90.0, 90.0 | 90.0, 109.2, 90.0 | 90.0, 90.0, 90.0 |
| Resolution range (Å)<sup>a</sup> | 83.9-2.70 (2.83-2.70) | 38.0-2.80 (2.89-2.80) | 77.2-2.80 (2.97-2.80) | 66.2-2.17 (2.24-2.17) | 41.8-1.86 (1.91-1.86) |
| R<sub>merge</sub><sup>ab</sup> | 0.107 (0.559) | 0.103 (0.685) | 0.267 (1.116) | 0.054 (0.508) | 0.077 (1.067) |
| R<sub>meas</sub><sup>ac</sup> | 0.115 (0.598) | 0.134 (0.891) | 0.299 (1.246) | 0.062 (0.594) | 0.096 (1.380) |
| CC<sub>1/2</sub><sup>ad</sup> | 0.996 (0.892) | 0.990 (0.580) | 0.977 (0.648) | 0.998 (0.852) | 0.998 (0.524) |
| Mean I/σ<sup>a</sup> | 12.9 (3.7) | 7.3 (1.3) | 6.2 (1.5) | 14.6 (2.3) | 11.2 (1.3) |
| Completeness (%)<sup>a</sup> | 99.6 (100.0) | 97.6 (97.7) | 99.4 (99.7) | 98.0 (91.6) | 99.5 (99.3) |
| Multiplicity<sup>a</sup> | 7.7 (8.1) | 2.2 (2.3) | 4.8 (5.0) | 4.5 (3.7) | 5.2 (5.1) |
| Wilson <B> (Å<sup>2</sup>) | 48.9 | 50.0 | 37.1 | 42.5 | 31.1 |
| **Refinement** |  |  |  |  |  |
| Resolution range (Å) | 64.1-2.70 | 38.0-2.80 | 60.3-2.80 | 66.2-2.17 | 39.0-1.86 |
| No. of reflections | 11513 | 49334 | 8975 | 21802 | 11247 |
| R<sub>work</sub>/R<sub>free</sub> | 0.2299/0.2635 | 0.2172/0.2751 | 0.2550/0.3121 | 0.1823/0.2293 | 0.2278/0.2636 |
| **Number of atoms** |  |  |  |  |  |
| Protein | 2030 | 12871 | 2307 | 2321 | 918 |
| Ligand/ion | 3 | 19 | 0 | 0 | 3 |
| Water | 43 | 232 | 35 | 196 | 121 |
| B-factors (Å<sup>2</sup>) | 55.6 | 51.3 | 40.1 | 49.5 | 38.8 |
| **Rmsd from ideal values** |  |  |  |  |  |
| Bond lengths (Å) | 0.006 | 0.013 | 0.004 | 0.005 | 0.021 |
| Bond angles (°) | 0.686 | 0.755 | 0.662 | 0.812 | 0.550 |
| **Ramachandran Plot** |  |  |  |  |  |
| Favoured region (%) | 92.5 | 91.7 | 87.3 | 95.3 | 98.3 |
| Allowed (%) | 100.0 | 99.7 | 97.9 | 100.0 | 100.0 |
| Outliers (%) | 0 | 0.3 | 2.1 | 0 | 0 |
| Rotamer outliers (%) | 0.9 | 0.3 | 1.2 | 0 | 0 |
| C-beta outliers | 0 | 1 | 0 | 0 | 0 |
| PDB ID code | 5MW5 | 5MWF | 5MV7 | 5MVX | 5MWB |

<sup>a</sup> Values in parentheses are for the highest resolution shell. <sup>b</sup> $R_{\text{merge}} = \sum(I_{hl} - \langle I_h \rangle) / \sum(I_{hl})$ where $\langle I_h \rangle$ is the mean intensity of unique reflection $h$ , summed over all reflections for each observed intensity $I_{hl}$ . <sup>c</sup> $R_{\text{meas}} = \sum(n/n - 1)^{1/2} (I_{hl} - \langle I_h \rangle) / \sum(I_{hl})$ where $n$ is the number of observations for unique reflection $h$ with mean intensity $\langle I_h \rangle$ , summed over all reflections for each observed intensity $I_{hl}$ . <sup>d</sup> CC<sub>1/2</sub> is the correlation coefficient on $\langle I \rangle$ between random halves of the dataset.

**Figure 1.**
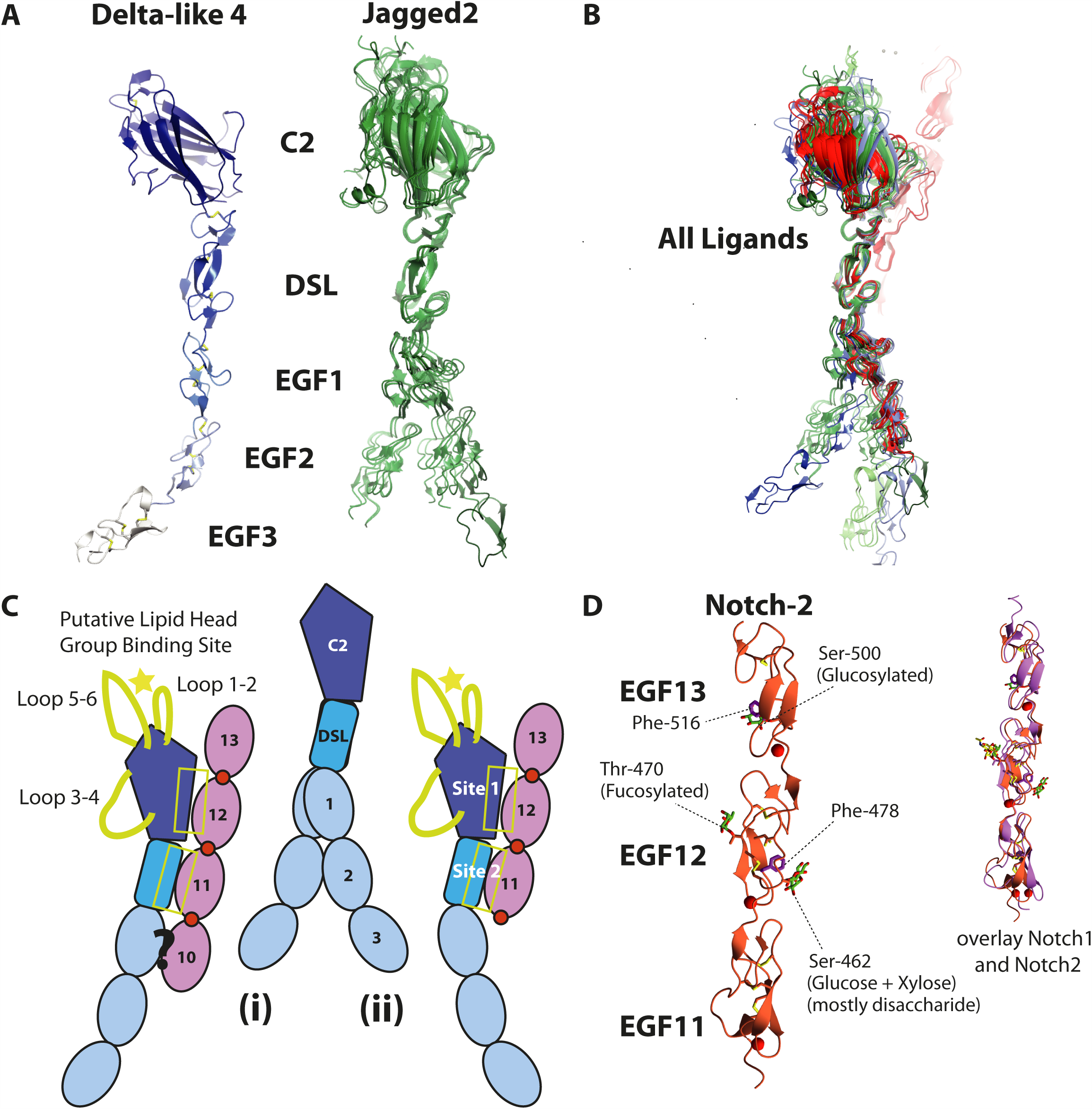
New structures of human Notch ligands and receptors. (**A**) Structures of human DLL4 (N-EGF3) and human Jagged2 (N-EGF2 and N-EGF3). Two crystal forms of Jagged2 (N-EGF2) (green) and one crystal form of Jagged2 (N-EGF3) (dark green) were solved, including a total of 7 crystallographically independent copies of N-EGF2, and 1 copy of N-EGF3 (N.b. Not all of EGF3 is visible in the electron density). All of these copies have been superposed on each other by alignment of the DSL domain, showing some flexibility in the hinge region between the C2 and DSL domains, and further flexibility in the hinge regions between the EGF domains. (**B**) Superposition of all of the known human Notch ligand structures (Jagged-1, PDB ID = 4CC1 (light green) (Chillakuri et al., 2013), DLL1, PDB ID = 4XBM (light blue) (Kershaw et al., 2015), DLL4 (blue), Jagged-2 (green)), and all of the DLL4 variant-Notch-1 complex structures (red) across the DSL domain, shows that these structures appear to fall into two groups (**C**). One group has an overall globally bent arrangement (**C(ii)**), which includes all of the DLL4 molecules bound to Notch-1 EGF11-13, and a second group with a straighter arrangement (**C(i)**). Binding of ligands to a Notch receptor in a native context likely requires the ligands to be in the straighter arrangement, as the bent arrangement is incompatible with binding to EGF10. (Weisshuhn et al., 2016). (**D**) Crystal structure of human Notch-2 (EGF11-13), and comparison with modified human Notch-1 EGF11-13 (PDB ID = 4D0E) (Taylor et al., 2014) (**B**). Phe-478 and Phe-516 are highlighted in Notch-2 as these appear to be shielded from the solvent by the glycans on Ser-462 and Ser-500 respectively.

### Comparison of the N-terminal C2 domains of human Notch ligands show differences in the lipid binding region

All of the C2 domains of the human Notch ligands superpose with RMSDs of between 1.1 and 1.5 Å. All have type II topology and are most similar to members of the PKC-C2 family (Corbalan-Garcia and Gomez-Fernandez, 2014) including Munc-13 and phosphoplipase A2 (cPLA2) (from DALI search). One distinct feature of the Notch ligand C2 domains is the presence of a long loop between strands 2 and 3, which forms the Notch EGF12 binding site 1. The stability of this loop is supported by a disulphide bond between the loop, and strand 2 (Luca et al., 2015). Despite strong overall structural conservation of the C2 domains there are major differences in the loops between strands 1 and 2, strands 3 and 4 and between strands 5 and 6. This region, at the extreme termini of the Notch ligands, is the putative lipid binding site (Figure 2A,B) and the differences in the loops of the ligand C2 domains most likely confer different lipid binding specificities. Even between rat and human DLL4 there are loop differences suggesting that these regions are optimized for the different mammalian physiologies (Figure S1).

**Figure 2.**
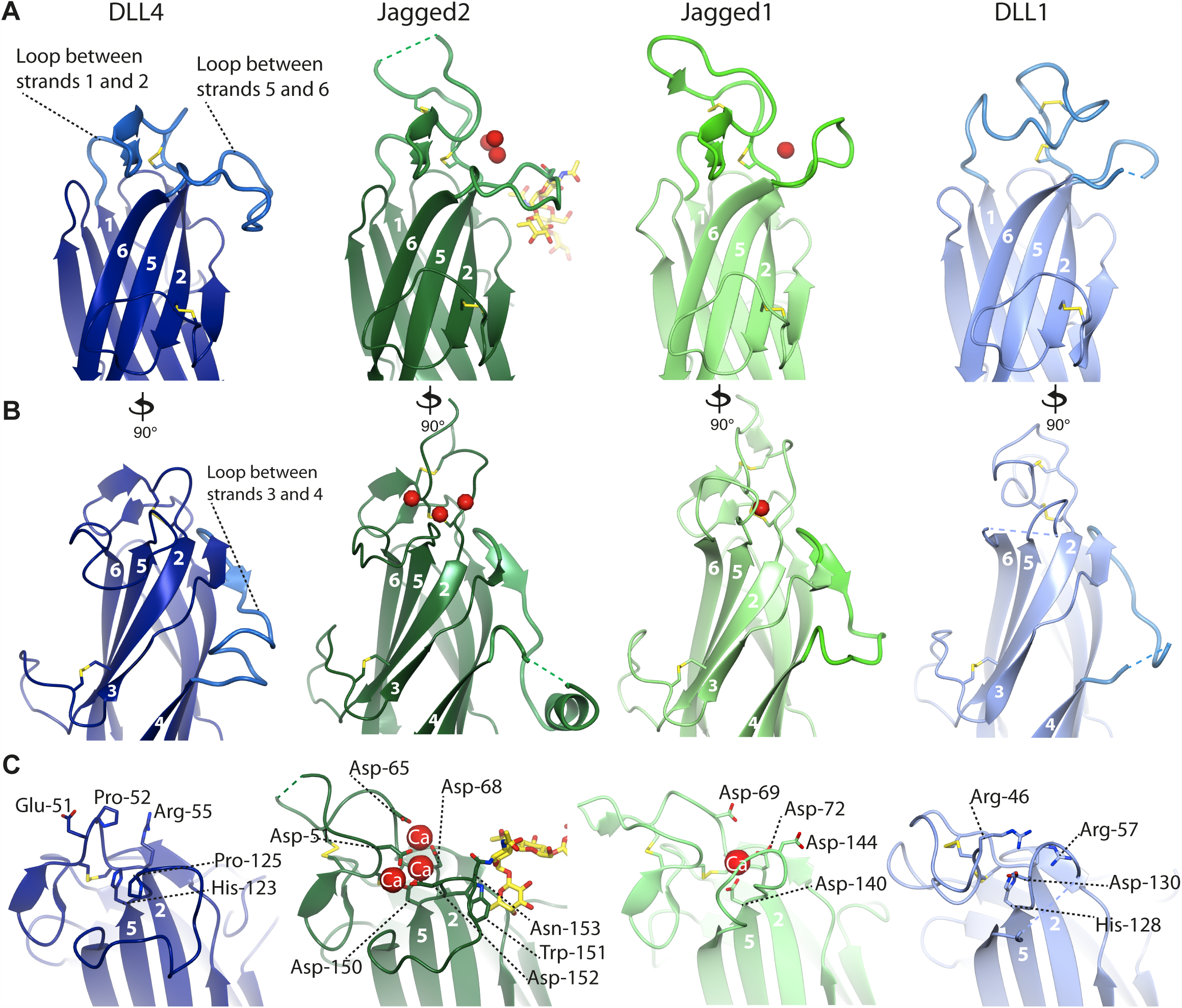
Comparison of the N-terminal C2 domains in the different Notch ligands. (**A**) The loops between strands 1 and 2 (Calcium binding region 1 (CBR1)), and between 5 and 6 (CBR3) of the Notch ligand C2 domains differ in length and conformation. (**B**) The loop between strands 3 and 4 (CBR2) is also different. (**C**) The Jagged1 and Jagged2 C2 domains bind calcium ions (shown in red), where as DLL4 and DLL1 do not (**C**); the aspartates involved in calcium binding are not conserved in DLL4 and DLL1. Both Jagged1 and Jagged2 contain N-glycosylation sites in the loop between strands 5 and 6. The α(1,6)-linked fucose on the Asn-153 N-glycosylation site in Jagged2 packs against Trp-151 side chain (shown). All of these differences at the putative lipid binding site likely reflects the different Notch ligand lipid binding specificity.

The C2 loops are ordered upon calcium binding in both Jagged1 (Chillakuri et al., 2013) and Jagged2 and contain the amino acid ligands for calcium binding although the number of calcium ions bound differs. Interestingly, the three calcium ions bound in the Jagged2 C2 domain are at equivalent positions to three of the five calcium ions bound in the Perforin C2 domain (Yagi et al., 2015). The loops are fully ordered in DLL4, and mostly ordered in DLL1 (Kershaw et al., 2015) despite the absence of calcium-binding sites in the Delta family (Figure 2A, B). Consistent with this, two of the key aspartate side chain calcium ligands conserved in Jagged1 and 2 are replaced with arginine and histidine residues (Arg-55 and His-123 in DLL4) (Figure 2C). All of the Notch ligands, irrespective of whether they have calcium-binding sites or not, contain few hydrophobic residues in calcium-binding region 1 (CBR1) or CBR3, and those that are present are not at the tip of the loops (tip being defined as closest to the membrane) (Figure 2). This suggests that the Notch ligand C2 domains are not deeply buried within the membrane upon binding and are therefore distinguishable from intracellular C2 domain proteins, and Perforin (Yagi et al., 2015).

### Human Notch-2 EGF11-13 structure is highly homologous to Notch 1

We have also solved the structure of human Notch-2 EGF11-13, which includes the ligand-binding region (EGF11-12) (Figure 1D and table 1). Human Notch-2 EGF11-13 has 67% sequence identity with human Notch-1, and superposition of the two structures shows that both adopt a very similar linear arrangement (RMSD = 1.55 Å) (Figure 1D). In our crystal structure, Notch-2 is O-fucosylated on Thr-470, and O-glucosylated on Ser-462 and Ser-500. In the earlier Notch1-DLL4 structure the O-fucose on the equivalent threonine residue in EGF12 is seen to be involved in direct intermolecular binding to the C2 domain (Luca et al., 2015; Taylor et al., 2014). In contrast the O-glucose modifications affecting specific serine residues in Notch-2 appear to stabilize intramolecular structure as observed previously for the Notch-1 complex (Luca et al., 2015): the disaccharide (glucose + xylose) on Ser-462 may stabilize Notch-2 EGF12 through interaction with Phe-478 (shown), shielding it from the solvent. Similarly the O-glucose on Ser-500 may stabilize EGF13 through interaction with Phe-516 (Figure 1D).

The availability of our new structures of hJagged2, hDLL4 and hNotch-2 allow modelling of the receptor-binding interface across all canonical ligands and Notch-1/Notch-2 and show that almost all of the key residues involved in ligand binding (Leu-468, Asp-469 and Ile-477 in site 1, Phe-436 and Arg-448 in site 2) are conserved in Notch-2. Receptor-binding residues in site 1 and 2 are also highly conserved across the ligands suggesting that other forms of regulation must be present to drive specific ligand-receptor pairing and signaling, rather than intrinsic differences in affinity between the core recognition elements of the various receptor-ligand pairs (Figure 3 & S1). This is supported by assaying different combinations of receptor/ligands which show similar levels of binding (Figure 4A). Fringe extension of O-fucosylated sites in the receptor is already known to change the responsiveness to ligand binding, but the variation in loop sequences in C2 domains of ligands offer another potential method to fine tune the affinity of the Notch complex and subsequent signaling capability.

**Figure 3.**
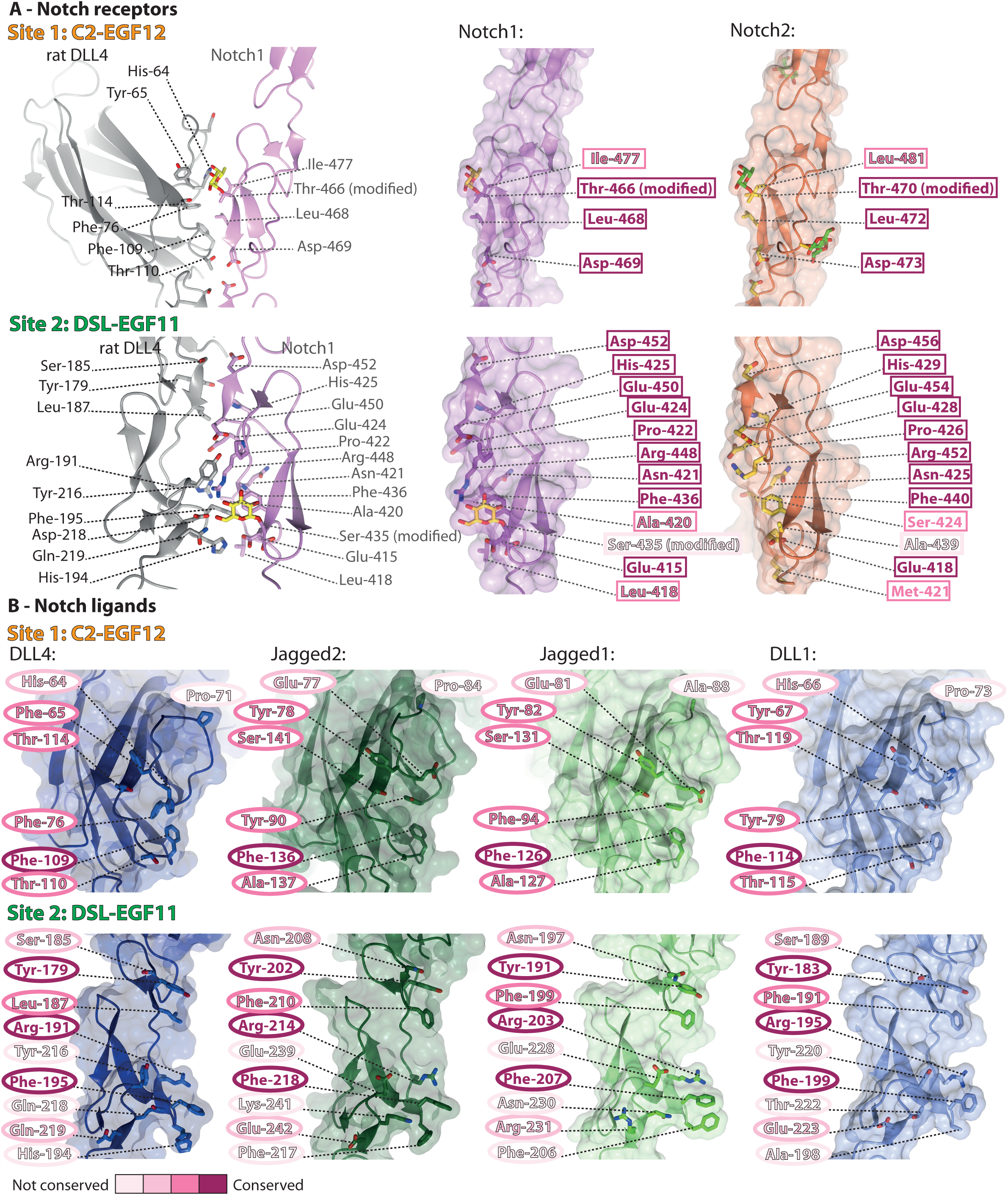
Comparison of the residues involved in complex formation in the different Notch receptors and Notch ligands. (**A**) Comparison of the residues involved in complex formation (Luca et al., 2015) in Notch-1 and Notch-2, and (**B**) in the different human ligands, at site 1 (C2:EGF12) and site 2 (DSL:EGF11). N.b. The complex structure between DLL4 and Notch-1 was of rat DLL4 (shown in panel A); human DLL4 is shown in panel B. Conservation of the residues involved in complex formation is indicated by background colour, highlighting the high conservation of the ligand binding sites in Notch-1 and Notch-2. There are a few residues in the receptor binding sites of the ligands that are absolutely conserved, with site 2 (DSL-EGF11) being more variable than site 1 (C2-EGF12).

**Figure 4.**
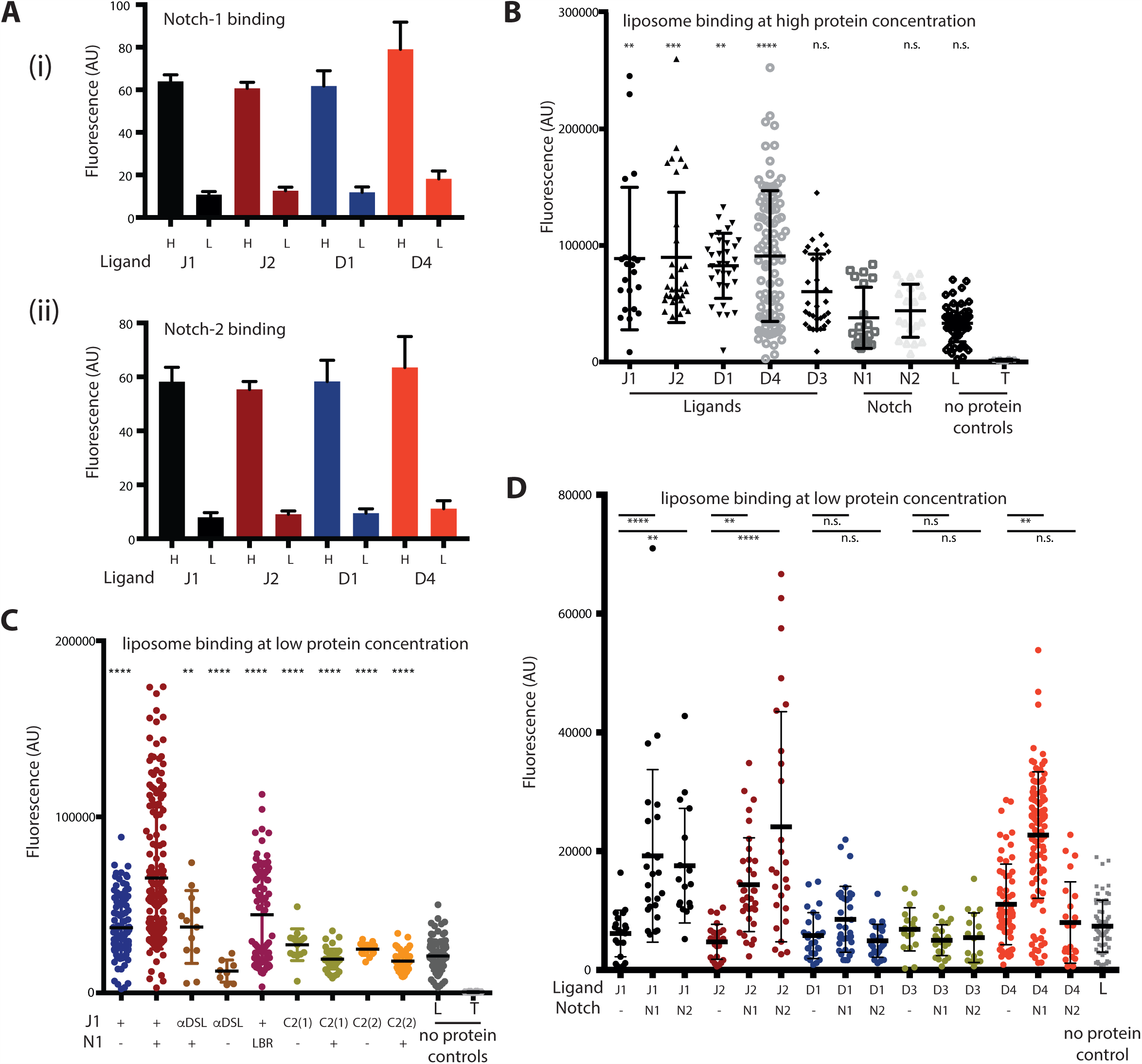
Functional coupling between liposome and Notch binding of Notch ligands. (**A**) All canonical human Notch ligands (Jagged1, Jagged2, DLL1 and DLL4) (N-EGF3) bind to both human Notch-1(i) and Notch-2 (ii) EGF11-13 to a similar extent as assessed by plate assay. Notch ligands were bound to nickel-coated plates before biotinylated pre-clustered Notch was added with NeutrAvidin conjugated HRP. Binding is shown at high (H) and low (L) protein concentrations, L indicating background. All components were purified from insect cells. (**B**) All canonical human Notch ligands (Jagged1 (J1), Jagged2 (J2), DLL1 (D1) and DLL4 (D4)) (N-EGF3) bound to fluorescently labeled liposomes (PC/PE/PS) (A) – DLL3 (N-EGF1), Notch-1 (EGF11-13) and Notch-2 (EGF11-13) did not bind. (**C**) At low concentrations of ligand i.e. below concentrations where liposome binding can be observed, addition of Notch-1 EGF11-13 stimulated binding of Jagged1 N-EGF3 to liposomes. This effect could be abolished by an antibody against the Notch-binding DSL domain of Jagged1 (α–DSL), or by substitution of residues critical for ligand binding to Notch (Leu468Ala) (LBR). Liposome binding was also abolished by substitutions that directly perturb the putative lipid binding site in the C2 domain of Jagged1 (Asp140Ala/Asp144Ala, C2(1) and Del1Del2Asp140Ala, C2(2)) (Chillakuri et al., 2013). (**D**) Addition of Notch-1 EGF11-13 stimulated binding of Jagged-1 and DLL4 N-EGF3 to liposomes, with Notch-2 having a similar effect on Jagged-2, but neither had an effect on DLL1 or DLL3 in terms of liposome binding. Statistical tests were performed as described in Supplementary Methods.

### Cross-talk between Notch and liposome binding

Purification of N-terminal fragments of all of the human Notch ligands, allowed us to assess lipid binding of these constructs. All of the canonical Notch ligands (Jagged1, Jagged2, DLL1 and DLL4) bound to liposomes consisting of a mixture of phosphatidylcholine/phosphatidylserine/phosphatidylethanolamine (PC/PS/PE), however DLL3 (non-canonical) did not show significant binding (Figure 4B). To investigate whether Notch ligands are able to bind to both lipids and Notch simultaneously we added Notch-1 EGF11-13 into our liposome assays. When working at low concentrations of ligand i.e. below concentrations where liposome binding can be observed, the addition of Notch-1 stimulated liposome binding to Jagged1 (Figure 4C). This stimulation is abolished when residues in the C2 domain involved in calcium binding were mutated (D140A/D144A, C2(1)), or when the two loops forming the putative lipid binding site in the C2 domain were substantially shortened (Del1Del2D140A, C2(2)). In addition, liposome binding was also abolished when Notch binding was inhibited by either inclusion of an antibody recognising the receptor-binding site of the DSL domain of Jagged1 (α-DSL), or use of a Notch-1 variant L468A, defective in ligand binding (LBR) (Figure 4C).

To investigate whether or not this stimulation of liposome binding is seen for all Notch ligands, we set up analogous liposome binding assays adding either Notch-1 or Notch-2 EGF11-13. Addition of Notch-1 to liposome assays enhanced binding of Jagged1, Jagged2 and DLL4 to liposomes, while Notch-2 enhanced binding to Jagged1, Jagged2 but not Dll4 (Figure 4D). No enhancement is seen with either Notch-1 or Notch-2 for DLL1 or DLL3 liposome binding. The enhancement of liposome binding seen for some of the ligands upon addition of Notch, may be due to Notch binding to and rigidifying the ligands, and thereby decreasing the loss of entropy upon binding to the lipids/membrane. Such coupling between the two binding events gives a mechanism to increase the affinity of the receptor/ligand complex and enhance signaling by a specific ligand in a particular physiological context and/or to affect selection by Notch of one ligand from a pool of ligands expressed on the cell surface, despite the apparently similar affinities of each Notch for all ligands. Thus the lipid composition of the cell membrane could act as a modulator of Notch signaling, in addition to O-glycosylation of the receptor.

### Human disease-associated mutations alter membrane but not Notch binding

Taken together our data strongly support a key role for the ligand C2 domains in Notch signaling independent of direct Notch engagement. Point mutations in *Jagged-1* have previously been linked with two diseases; Alagille Syndrome (Penton et al., 2012) and Extrahepatic Biliary Atresia (Kohsaka et al., 2002). Alagille is a more severe multi-system disease and we have previously demonstrated that the amino acid substitutions linked to this disease result in mis-folded protein being retained within the cell and hence lead to haplo-insufficiency of Jagged-1 (Chillakuri et al., 2013). The *Jagged-1* point mutations in EHBA affect only one organ system and lead to an abnormal or absent extrahepatic bile duct implying a more subtle effect on Notch-signalling. We therefore generated recombinant Jagged1 variant proteins bearing three of the individual amino acid substitutions associated with EHBA – Asn53Asp and Lys65Met within the apical loops of the C2 domain and Arg203Lys within the Notch-binding interface of the DSL (Figure 5A). All these variant forms could be purified as recombinant proteins unlike those previously studied Alagille variants, suggesting that haploinsufficiency of Jagged1 does not cause EHBA. All variants reduced activation in a cellular assay of Notch-signalling activity to a similar level to control substitutions (Phe207Ala within the Notch-binding site or Asp140Ala/Asp144Ala within the C2 domain). (Figure 5B). To define the mechanisms leading to reduction in activity we next tested the variant proteins to see if either membrane or Notch recognition were altered. As predicted, the disease-causing variant within the Notch-binding site abrogated Notch binding (Figure 5C) but left liposome binding intact (Figure 5D). Conversely, both disease-causing C2 domain variants left Notch binding unaltered (Figure 5C), but significantly reduced liposome binding (Figure 5D). All three disease causing variants therefore did not show the enhancement of Notch-binding in the presence of liposomes seen for WT Jagged1 (Figure 5E). These data strongly suggest that EHBA is caused by a failure to form a ternary complex comprising membrane, Notch and ligand, which in the cases studied here can be due to a reduction in membrane recognition by the C2 domain (Asn53Asp, Lys65Met) or a direct effect on Notch recognition mediated by the DSL domain (Arg203Lys). These data therefore strongly support the idea that two recognition events are critical for efficient Notch-signalling in some physiological contexts, such as bile duct development.

**Figure 5.**
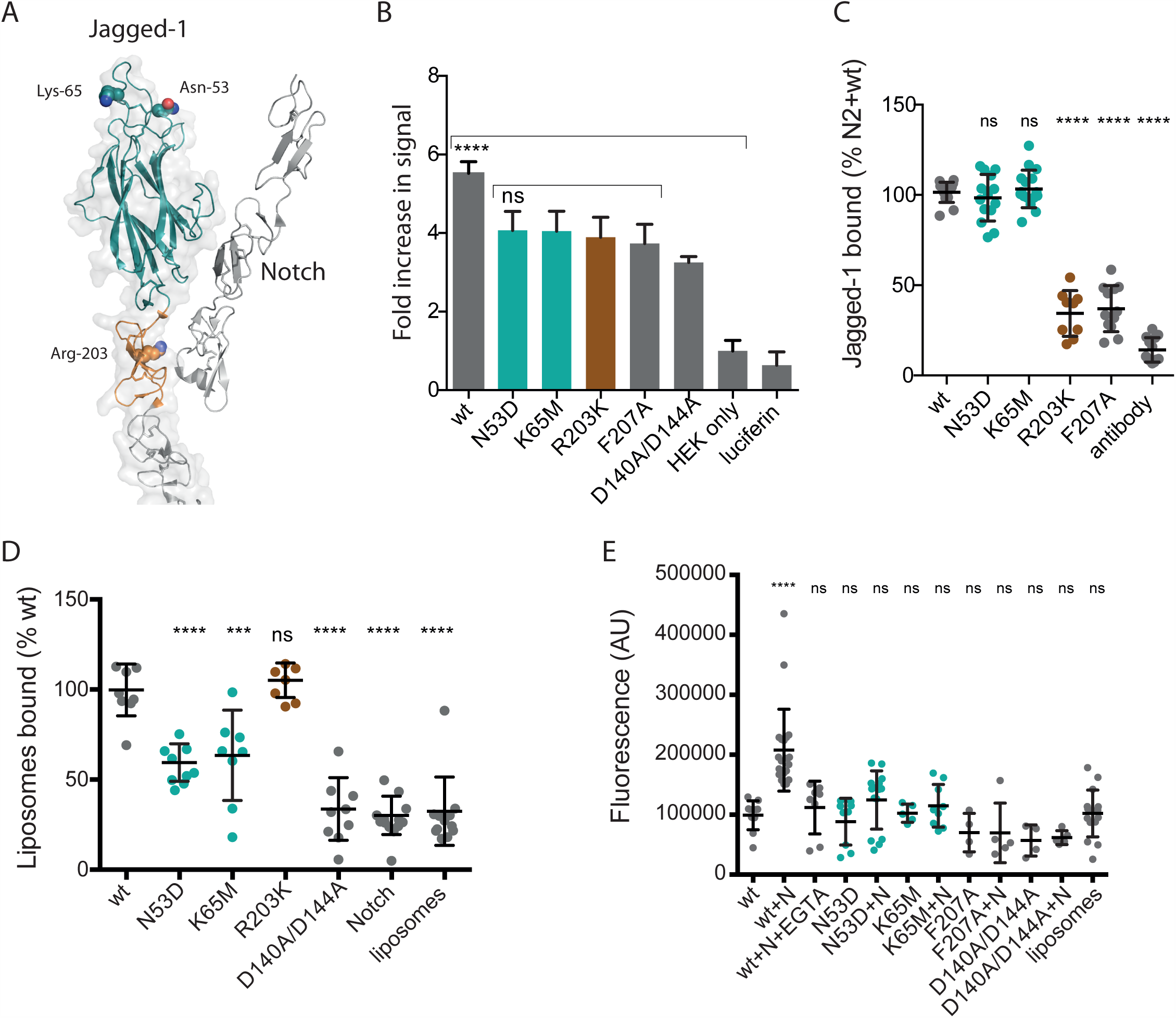
Disease-causing substitutions affecting C2 loops selectively alter membrane but not Notch binding. (**A**) C2 domain of Jagged1 showing position of extrahepatic biliary atresia (EHBA) causing substitutions Asn53Asp and Lys65Met in loop regions, the position of Arg203Lys in DSL Notch binding site also associated with EHBA is shown. (**B**) C2 EHBA variants reduce Notch-1 and Notch-2 activation and (**C**) C2 EHBA variants do not affect Notch binding, unlike Arg203Lys but (**D,E**) liposome binding is reduced and the Notch boosting effect is lost. Statistical tests were performed as described in Supplementary Methods.

Since composition of membrane can vary with cell type and stage of development, future studies will investigate the *in vivo* importance of the ternary complex for Notch signaling utilizing CRISPR-cas genome editing approaches in combination with structure-informed mutagenesis of C2 domain loops from different Notch ligands.

## Methods

### Protein expression and production

Notch ligand and receptor constructs were recombinantly expressed in S2 insect cells (Expres2ion^®^ Biotechnologies, Denmark) as C-terminally His-tagged fusion proteins. Media containing recombinantly expressed protein was filtered and loaded onto a complete His-tag Purification Column (Roche Diagnostics, UK), for purification via His-tag. Following washing with 50 mM Tris pH 9.0, 5 mM Imidazole pH 8.0, 200 mM NaCl, 1 mM CaCl_2_, proteins were eluted with buffer containing 250 mM Imidazole pH 8.0. Following overnight dialysis proteins were further purified by size-exclusion chromatography (SEC) using a Superdex S200 (ligands) or S75 (receptors) preparative column in 20 mM Tris pH 7.5, 200 mM NaCl, 1 mM CaCl_2_.

### Structure determination of extracellular fragments of human Notch ligands and the Notch-2 receptor

DLL4 N-EGF3 was crystallised at 3 mg/ml in 0.1 M MES pH 6.5, 12% (w/v) PEG-20K at a 3:1 protein:precipitant ratio. Crystals were cryoprotected with 35% (v/v) glycerol, and data were collected to 2.2 Å (Table 1). The crystal belonged to space group C2, with one molecule in the asymmetric unit. The structure was solved by molecular replacement using the C2 domain of Jagged1 (PDB ID = 4CC1) (Chillakuri et al., 2013) using *Phaser* (McCoy et al., 2007) within *CCP4* (Winn et al., 2011), before the remaining domains were found also using Jagged1 as a search model. The structure was built using the automatic model building software *Buccaneer* (Cowtan, 2006). All structures were refined using *Coot* (Emsley et al., 2010), *Refmac* (Murshudov et al., 1997) and *Phenix refine* (Afonine et al., 2012).

Jagged2 N-EGF3 was crystallised at 2.3 mg/ml in the presence of 10 mM BaCl_2_ in 0.1 M sodium citrate pH 5.3, 20% (w/v) PEG-5000 MME at a 3:1 protein:precipitant ratio. Crystals were cryoprotected with 30% (v/v) ethylene glycol, and data were collected to 2.8 Å (Table 1). The crystal belonged to space group P2_1_2_1_2_1_, with one molecule in the asymmetric unit. The structure was solved by molecular replacement using the C2 domain of Jagged1 (PDB ID = 4CC1) (Chillakuri et al., 2013) using *Phaser* (McCoy et al., 2007), before the remaining domains of Jagged1 N-EGF3 were placed sequentially into the electron density with iterative rounds of rigid body and restrained refinement in *Refmac* (Murshudov et al., 1997). Barium was not visible in the electron density maps, and neither were the loops at the tip of the C2 domain.

Jagged2 N-EGF2 was crystallised at 3.4 mg/ml in the presence of 10 mM CaCl_2_ in 0.05 M potassium dihydrogen phosphate, 20% (w/v) PEG-8000 at a 3:1 protein:precipitant ratio. Crystals were cryoprotected with 20% (v/v) glycerol, and data were collected to 2.7 Å (Table 1). The crystal belonged to space group P2_1_2_1_2_1_ (similar to the apo N-EGF3 crystal form), with one molecule in the asymmetric unit. The structure was solved by molecular replacement using the N-EGF2 portion of the apo N-EGF3 model using *Phaser* (McCoy et al., 2007). Three calcium ions and most of the residues in the loops of the C2 domain are visible in the electron density.

Jagged2 N-EGF2 also crystallised at 3.3 mg/ml in the presence of 20 mM CaCl_2_ in 0.1 M sodium cacodylate pH 6.5, 0.2 M ammonium sulphate, 30% (w/v) PEG-8000 at a 1:1 protein:precipitant ratio. Crystals were cryoprotected with 20% (v/v) ethylene glycol, and data were collected to 2.8 Å (Table 1). The crystal belonged to space group C2, with six molecules in the asymmetric unit. The structure was solved by molecular replacement using the N-EGF1 portion of the above N-EGF2 structure model using *Phaser* (McCoy et al., 2007), before the EGF2 domains were placed manually into the electron density. Three calcium ions are bound to the C2 domain, with the loops of the C2 domain mostly visible (excluding three residues in the loop between strands 1 and 2). Asn-153 is N-glycosylated with density representing the first four sugar moieties visible (until the β(1-4)-linked mannose).

Notch2 EGF11-13 crystallised at 20 mg/ml in the presence of 10 mM CaCl_2_ in 0.1 M sodium cacodylate pH 6.5, 0.2 M sodium acetate, 30% (w/v) PEG-8000 at a 3:1 protein:precipitant ratio. Crystals were cryoprotected with 15% (v/v) ethylene glycol, and data were collected to 1.9 Å (Table 1). The crystal belonged to space group P2_1_2_1_2_1_, with one molecule in the asymmetric unit. The structure was solved by molecular replacement using the individual EGF domains of Notch1 (PDB ID = 2VJ3) (Cordle et al., 2008). EGF12 was found first, before EGF11 and EGF13. Density representing *O*-glucose on Ser-462 and Ser-500, and *O*-fucose on Thr-470 was clearly visible. Density representing xylose linked to the *O*-glucose on Ser-462 was also visible.

### Determination of Apparent Binding Affinities by Plate assay

Pierce 96-well nickel coated plates (Thermo Scientific) were coated with Notch ligands (5 μg/ml) (N-EGF3 constructs) in 20 mM HEPES pH 7.4 containing 200 mM NaCl (HBS). After incubation at 4°C overnight, the plate was washed and blocked for 1 hour at room temperature with 4% (w/v) Bovine Serum Albumin in the same buffer. Pre-clustered Notch constructs (0–300 nM) were added to the ligand-coated plates and incubated for 1.5 hour at room temperature. After washing four times with HBS containing 5 mM CaCl_2_ and 0.05% (v/v) Tween 20, followed by two washes in the absence of Tween 20, the plate was developed with 2,2′-Azino-bis(3-ethylbenzothiazoline-6-sulfonic acid) diammonium salt (Sigma-Aldrich). Absorbance was measured with a PHERAstar FS microplate reader (BMG LABTECH).

### Liposome binding assays

Notch ligands (e.g. Jagged1 N-EGF3) were coated at 800 nM/ 200 nM in a 40 μl volume onto a nickel coated plate (Pierce Nickel Coated Plates, black, 96-well, #15342) in 20 mM HEPES pH 7.4, 200 mM NaCl through incubation overnight at 4°C. The former concentration was used for testing liposome binding of the Notch ligands, and the latter for incorporating hNotch1 EGF11-13 into the assays.

Wells were washed twice with 20 mM HEPES pH 7.4, 200 mM NaCl before blocking for ˜3 hours with 0.1% (w/v) gelatin in the same buffer. Following incubation, wells were washed twice more with buffer, with the second wash buffer also containing 5 mM CaCl_2_.

To appropriate wells, 8 μl of anti-Jagged1 DSL domain Ab supernatant was added.

Liposomes were prepared as described in Chillakuri *et al.* (Chillakuri et al., 2013). To analyse the effect of adding Notch1 EGF11-13 on liposome binding, Notch1 EGF11-13 was added at a 4:5 liposome:protein volume ratio, with liposomes diluted 1 in 20 in 20 mM HEPES pH 7.4, 200 mM NaCl, 2 mM CaCl_2_, and Notch1 at 800 nM in the same buffer.

The liposome:protein mix was incubated on a rotating shaker for 1 hour, before washing twice in 20 mM HEPES pH 7.4, 200 mM NaCl, 5 mM CaCl_2_. Liposomes were solubilised in 40 μl of 0.3% Triton X-100 for at least 1 hour before the fluorescence intensity was measured in 96-well plate reader format using PHERA Star BMG Labtech (wavelengths: excitation 485nm, emission 520nm).

### Assay of EHBA variant proteins

Ligands proteins were produced as Fc fusions in human embryonic kidney 293T (HEK293T) cells using a transient transfection system (Aricescu et al., 2006) and purified as described in Chillakuri *et al.*, 2013 (Chillakuri et al., 2013). Notch1/2 EGF11-13 constructs were produced in S2 system and purified as above. Notch activation assays were performed as in Chillakuri *et al.*, 2013 with different reporter cell lines. Notch-2 and Notch-1 cell lines were a kind gift from R. Kopan. These cell lines constitutively express Notch-1 and 2, respectively, thus assays were performed without addition of doxycycline (Liu et al., 2013). Notch binding to EHBA variants was performed according to the following: Briefly 200 ng (50 μl volume) of monomeric Notch-1/2 EGF11-13 construct in 20 mM HEPES pH 7.4, 200 mM NaCl buffer was immobilized on a transparent 96 well Maxisorp immunoplate overnight at 4°C. The wells were then washed three times with 200 μl buffer and blocked with 200 μl of blocking buffer: 20 mM HEPES pH 7.4, 200 mM NaCl, 3% milk, 0.1% gelatin, 5 mMCaCl_2_. After 90 minutes (25°C), wells were washed and a 50 μl solution of 200 nM dimeric ligands constructs was added. 90 minutes later, wells were washed and incubated with a 1:2000 dilution of mouse-anti-human IgG Fc antibody conjugated to HRP for 60 minutes. After washing, 100 μl of 2,2′- Azino-bis(3-ethylbenzothiazoline-6-sulfonic acid) diammonium salt (Sigma-Aldrich) substrate solution was added to each well. The absorbance of the plate was read at a wavelength of 415 nm using a Pherastar plate-reader.

Liposome binding experiments were performed as above except black plates were coated with 40 μl of 800 nM ligands in 20 mM HEPES pH 7.4, 200mM NaCl. To determine if Notch increased liposome binding to EHBA variants 40 μl of 200 nM dimeric ligands were coated overnight at 4°C onto a black plate in 20 mM HEPES pH 7.4, 200 mM NaCl and the assay performed as described above.

### Statistical analyses

All data were analysed with the Prism 6 software (GraphPad, San Diego, CA, USA). Comparisons between two groups were performed with two-tailed unpaired *t*-test. Statistical differences among various groups were assessed with ordinary one-way Anova by comparison to the mean of a control column. Values are presented together with mean±SD.

## Acknowledgements

RJS & BK were funded by a Medical Research Council Grant MR/L001187/1 & CM by Wellcome Trust Grant 097928 to PAH & SML. SML is supported by a Wellcome Trust Investigator Award 100298. PW is supported by a grant awarded by the EPA Cephalosporin Fund. We thank Diamond Light Source for access to beamlines I02, I03, I04 & I04-1 (MX12346) that contributed to the results presented here. All protein structures and X-ray data have been deposited in the Protein Data Bank with the identifiers shown in Table 1.

---

**Figure S1 - Structural comparison of human DLL4 ligand with rat DLL4**

Superposition of human DLL4 (N-EGF3) (**A**) with rat DLL4_SLP_(N-EGF1) (grey) –Notch-1(EGF11-13) (pink) complex structure (PDB ID = 4XL1) (Luca et al., 2015) across the DSL domain, highlights the greater angle between the C2 and DSL domains in the apo ligand structure (**B**). There are also differences in the loops between strands 3 and 4 (CBR2), and strands 5 and 6 (CBR3) in the C2 domain between rat and human DLL4 (**C**). The Consurf server (consurf.tau.ac.il) was used to create a surface representation of the evolutionary conservation of residues in DLL4 based on an alignment from zebrafish to humans using ClustalO (data not shown). This highlights that CBR2 and CBR3 are the most variable regions of the structure (**C**).

